# Pancreatic cancer mutationscape: revealing the link between modular restructuring and intervention efficacy amidst common mutations

**DOI:** 10.1101/2024.01.27.577546

**Authors:** Daniel Plaugher, David Murrugarra

## Abstract

There is increasing evidence that biological systems are modular in both structure and function. Complex biological signaling networks such as gene regulatory networks (GRNs) are proving to be composed of subcategories that are interconnected and hierarchically ranked. These networks contain highly dynamic processes that ultimately dictate cellular function over time, as well as influence phenotypic fate transitions. In this work, we use a stochastic multicellular signaling network of pancreatic cancer (PC) to show that the variance in topological rankings of the most phenotypically influential modules implies a strong relationship between structure and function. We further show that induction of mutations alters the modular structure, which analogously influences the aggression and controllability of the disease *in silico*. We finally present evidence that the impact and location of mutations with respect to PC modular structure directly corresponds to the efficacy of single agent treatments *in silico*, because topologically deep mutations require deep targets for control.

## 1 Introduction

Biologically, regulatory networks and protein-protein interaction networks are typically thought to be densely connected sub-regions of an overall sparse system [1]. Natural cellular functions such as signal transmission are carried out by so-called modules that are discrete entities with separable functionality from other modules [2]. For example, the ribosome is a module that is responsible for synthesizing proteins that is spatially isolated. A similar isolation is seen with the proteasome. Whereas, signaling systems through chemokines would be extended modules that are isolated through the binding of chemical signals to receptor proteins. These isolating features allow cells to achieve various objectives with minimal influence from cross-talk [2]. Yet, their connectivity allows complex guidance signals from one another.

More often, *in silico* models are being implemented in cancer research for the discovery of general principles and novel hypotheses that can guide the development of new treatments. Despite their potential, concrete examples of predictive models of cancer progression remain scarce. One reason is that most models have focused on single–cell type dynamics, ignoring the interactions between cancer cells and their local tumor microenvironment (TME). There have been a number of models that were used to study gene regulation at the single-cell scale, such as macrophage differentiation [3, 4, 5], T cell exhaustion [6], differentiation and plasticity of T helper cells [7, 8], and regulation of key genes in different tumor types [9, 10], including pancreatic cancers [11].

These models are all great steps towards control-based treatment optimization, but it has been demonstrated that that the TME has a critical effect on the behaviour of cancer cells [12]. Ignoring the effect of cells and signals of the TME can generate confounding conclusions. For example, it was shown that in non-small cell lung cancer, the microenvironments of squamous tumors and adenocarcinomas are marked by differing recruitment of neutrophils and macrophages, respectively. *Ex vivo* experiments revealed the importance of the TME as a whole, especially when considering immunotherapy enhancement [13]. A similar observation has shown that removing pancreatic stellate cells from the TME *in silico* led to differing long-term outcomes because they form a protective layer around tumor cells [11].

To study the interplay of cancer cells with components of the TME, modelers developed multicellular models including cancer, stromal, immune, cytokines, and growth factors [14]. These models are typically multiscale integrating interactions at different scales, making it possible to simulate clinically relevant spatiotemporal scales and at the same time simulate the effect of molecular drugs on tumor progression[15, 16, 17, 18, 19, 20]. The high complexity of these models generates challenges for model validation such as the need to estimate too many model parameters.

While a a multi-scale model would likely provide more realistic simulations, state of the art control techniques require a network model specification such as Boolean networks (or their generalizations) to find optimal therapeutic interventions [21]. Implementation of Boolean networks (BNs) provides a coarse grained description of signaling cascades without the need for tedious parameter fitting and can be simulated through stochastic discrete dynamical systems (SDDS [22]) to streamline the modeling process and increase efficiency. These models have a well-studied and effective track record for capturing various biological system dynamics [23].

Pancreatic cancer is among the most lethal types of malignancies largely due to its difficulty to detect. The pancreas is located deep within body, and a standard doctor’s exam will likely not reveal a tumor. Additionally, there is an absence of detecting and imaging techniques for early stage tumors. While PC only accounts for 3% of estimated new cases, it is fourth highest cause of cancer-related death in the United States [24], and there is only a 3% five year survival rate among its late stage patients (approximately 82% in Stages 3 and 4) [25, 26]. PC is widely known for its resistance to most traditional therapy protocols. According to the Pancreatic Cancer Action Network (PanCAN), most patients receive fluorouracil (5-FU) or a gemcitabine based treatments that are anti-metabolites targeting thymidylate synthase and ribonucleotide reductase, respectively [26].

In a prior work, we presented a multicellular model of pancreatic cancer (PC) based on a stochastic Boolean network approximation (Figure 10 in Section 10), and we used control strategies that direct the system from a diseased state to a healthy state by targeting and disrupting specific signaling pathways. The model consists of pancreatic cancer cells (PCCs), pancreatic stellate cells (PSCs), cytokine molecules diffusing in the local microenvironment, and internal gene regulations for both cell types [11]. We then used the PC model to study the impact of four common mutations: KRAS, TP53, SMAD4, and CDKN2A [14, 27, 28]. Throughout this writing, we will often denote PCC components with a subscript *c* and PSC components with subscript *s*.

Using our PC model as a case-study, readers will find the following: Figure 1 shows a workflow of our process for defining and analyzing modules, Section 2 defines the methods we have implemented, followed by a summary of the PC model dynamics and target efficacy in Section 3. We show that the modularity of GRNs is vulnerable to mutations (Section 4. 1), analogous perturbations to long-term dynamical outcomes occur that influences aggression and controllability (Sections 4. 1 and 4. 2), variance in topological rankings of the most phenotypically influential modules implies a strong relationship between structure and function (Section 4. 2), and we finally present evidence that the impact and location of mutations with respect to modular structure directly corresponds to the efficacy of single agent treatments *in silico* (Section 4. 3).

**Figure 1:**
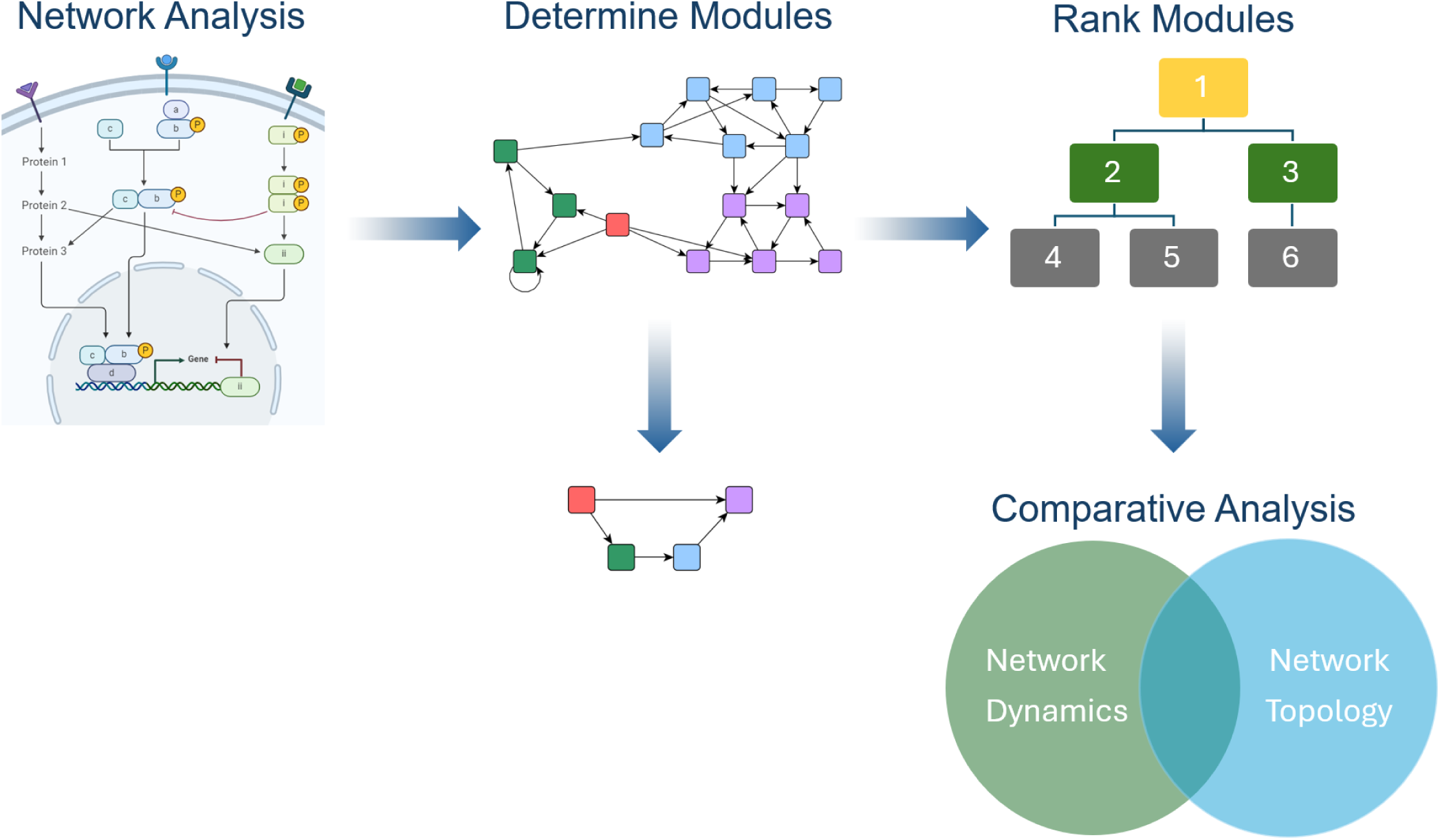
Modularity workflow. The workflow within this project begins with formal network analysis (previously conducted on a published PC model [11, 23, 29]). We then determine modules and reduce to non-trivial modules, rank the modules, and perform comparative analysis to understand the connection between dynamics and topology.

## 2 Methods

### 2. 1 Network modularity

Systems biology can often build complicated structures from simpler building blocks, even though these simple blocks (i.e. modules) traditionally are not clearly defined. The concept of modularity detailed in [1] gives a structural decomposition that induces an analogous decomposition of the dynamics of the network, in the sense that one can recover the dynamics of the entire network by using the dynamics of the modules. First, we note that the wiring diagram of a Boolean network is either strongly connected or is composed of strongly connected components (SCCs). Using this decomposition, we will define a *module* of a BN as a subnetwork that is itself a BN with external parameters in the subset of variables that specifies an SCC (example below). Naturally, this decomposition imposes a hierarchy among the modules that can be used for the purpose of control. In this work, we show one way to rank the modules to study their relevance with respect to effectiveness and aggressiveness of treatment.

More precisely, for a Boolean network *F* and subset of its variables *S*, we define a *subnetwork* of *F* as the restriction of *F* to *S*, denoted *F |_S_* = (*f_k_*1 *, · · ·, f_k_m*) where *x_k_i ∈ S* for *i* = 1*, · · ·, m*. We note that *f_k_i* might contain inputs that are not part of *S* (e.g,. when *x_k_i* is regulated by variables that are not in *S*). Therefore, *F |_S_* is a BN with external parameters. For the Boolean network *F* with wiring diagram *W*, let *W*_1_*, …, W_m_* be the SCCs of *W* with variables *S_i_*. The *modules* of *F* are *F |_S_i*, and setting *W_i_ −→ W_j_* where there exists a node from *W_i_* to *W_j_* gives a directed acyclic graph *C* = *{*(*i, j*)*|W_i_ −→ W_j_}* [1].

The dynamics of the state-space for Boolean network *F* are denoted as *D*(*F*), which is the collection of attractors. Further, if *F* is decomposable (say into subnetworks *H* and *G*), then we can write *F* = *H* ⋊*_P_ G* which is called the *coupling* of *H* and *G* by scheme *P*. In the case where the dynamics of *G* are dependent on *H*, we call *G non-autonomous* denoted as *G̅*. Then we adopt the following notation: let *A* = *A*_1_ *⊕ A*_2_ be an attractor of *F* with *A*_1_ *∈ D*(*H*) and *A*_2_ *∈ D*(*G^A^*1) [1].

Lastly, a set of controls *u* stabilize a BN to attractor *A* if the only remaining attractor after inducing *u* is *A*. The decomposition strategy can be used to obtain controls for each module, that can then combine to control the entire system. That is, given a decomposable network *F* = *F*_1_ ⋊*_P_ F*_2_ and an attractor *A* = *A*_1_ *⊕ A*_2_ with *A*_1_ *∈ D*(*F*_1_) and *A*_2_ *∈ D*(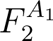), assume *u*_1_ is a control that stabilizes *F*_1_ in *A*_1_, and *u*_2_ is a control that stabilizes 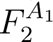 in *A*_2_. Then *u* = (*u*_1_*, u*_2_) is a control that stabilizes *F* in *A* given that either *A*_1_ or *A*_2_ are steady states [1].

For an example, consider the network in Figure 2a, which can be written as

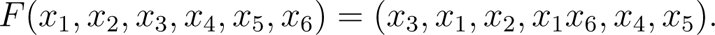

**Figure 2:**
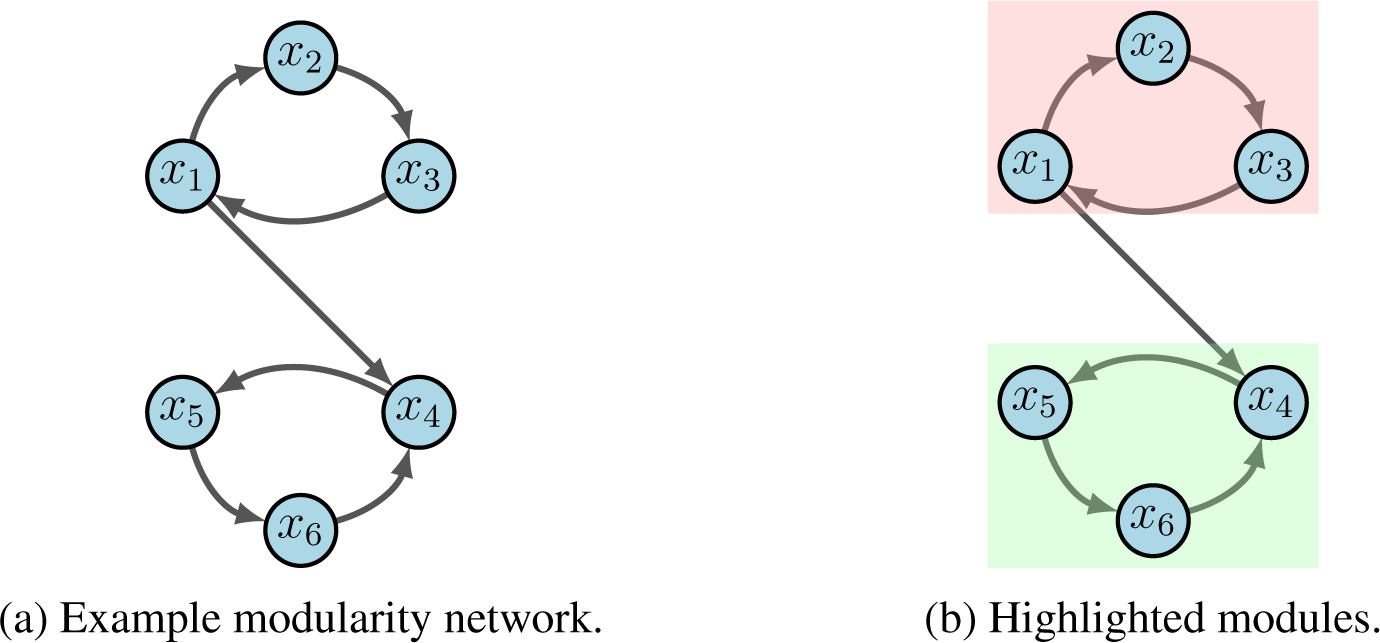
Modularity example.

Subnetworks are defined according to the dependencies of variables encoded by the wiring diagram [1]. For example, the subnetwork *F |_{x_*4*,x*5*,x*6*}* = (*x*_1_*x*_6_*, x*_4_*, x*_5_) is the restriction of *F* to *{x*_4_*, x*_5_*, x*_6_*}* with external parameter *x*_1_. From the given wiring diagram, we derive two SCCs where Module 1 (red in 2b) flows into Module 2 (green in 2b). That is, *F* = *F*_1_ ⋊ *F*_2_ with

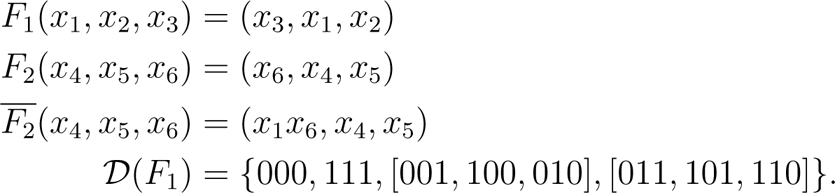

Suppose we aim to stabilize the system into *y* = 000000. First we see that either *x*_1_ = 0, *x*_2_ = 0 or *x*_3_ = 0 stabilize Module 1 (i.e. *F*_1_) to *A*_1_ = 000 by applying the Feedback Vertex Set method [30, 31]. Likewise, *x*_4_ = 0, *x*_5_ = 0 or *x*_6_ = 0 stabilize Module 2 (i.e. 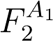) to *A*_2_ = 000. Thus, we conclude that *u* = (*x*_1_ = 0*, x*_6_ = 0) achieves the desired result.

### 2. 2 Topological sorting

To rank the modules of a Boolean network, we first formed the *condensation graph C* of its wiring diagram *W* which is obtained by contracting each strongly connected component into a single node (as in the middle panel of Figure 1). Thus, the condensation graph is a directed, acyclic graph whose nodes represent the modules of the original graph *W*. To obtain the modules and their components we used the MATLAB [32] function ‘condensation’ that returns the condensation graph *C*. Another MATLAB [32] function called ‘conncomp’ bins nodes according to their corresponding strongly connected component. Two nodes belong to the same strong component only if there is a path connecting them in both directions. Condensation determines the nodes and edges in *C* by the components and connectivity in *W* such that: *C* contains a node for each strongly connected component in *W*, and *C* contains an edge between node *I* and node *J* if there is an edge from any node in component *I* to any node in component *J* of *W*.

We proceeded to then order the modules using the topological ordering of an acyclic graph, which is an ordering of the nodes in the graph such that each node appears before its successors. We use an implementation of the topological sorting algorithm called ‘toposort’ from MATLAB [32]. The algorithm is based on a depth-first search, where a node is added to the beginning of the list after considering all of its descendants and returns a new order of nodes such that *i < j* for every edge (ORDER(*i*), ORDER(*j*)) in the original graph *W*. For example, consider a directed graph whose nodes represent the courses one must take in school and whose edges represent dependencies that certain courses must be completed before others. For such a graph, the topological sorting of the graph nodes produces a valid sequence in which the tasks could be performed [32]. Finally, we ranked the modules based on the percentile scores (i.e., rank module *k* out of *m* modules).

### 2. 3 Stochastic discrete dynamical systems

Synchronous updating schedules produce deterministic dynamics, where all nodes are updated simultaneously (i.e. in sync). The stochastic discrete dynamical systems (SDDS) framework developed by [22] incorporates Markov chain tools to study long-term dynamics of Boolean networks. By definition, an SDDS on the variables *x*_1_*, x*_2_*, …, x_n_* is a collection of *n* triples denoted 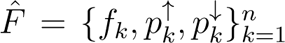, where for *k* = 1*, …, n*,

- *f_k_*: *{*0, 1*}^n^ → {*0, 1*}* is the update function for *x_k_*
- 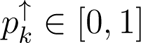 is the activation propensity
- 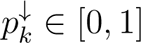 is the deactivation propensity

Consider the state-space *S*, consisting of all possible states of the system. If *x* = (*x*_1_*, …, x_n_*) *∈ S* and *y* = (*y*_1_*, …, y_n_*) *∈ S*, then the probability of transitioning from *x* to *y* is

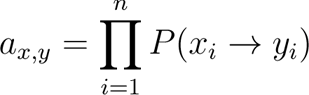

where

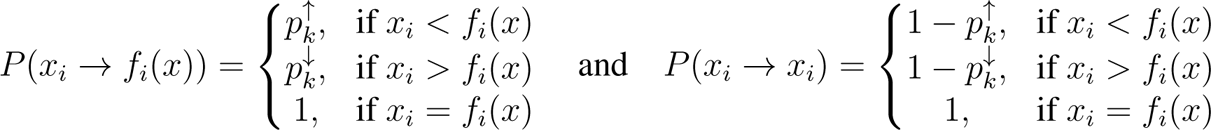

Here we assume that *P* (*x_i_ → y_i_*) = 0 for any *y_i_ ∈/{x_i_, f_i_*(*x*)*}*. When propensities are set to *p* = 1, we have a traditional BN [9]. With this framework, we built a simulator that takes random initial states as inputs and then tracks the trajectory of each node through time. Long-term phenotype expression probabilities can then be estimated, as well as network dynamics with (and without) controls.

## 3 Model dynamics and target efficacy

Prior work first completed a rigorous dynamical and network cascade analysis of the original non-mutant model [11]. We also identified numerous targets using various techniques for phenotype control including computational algebra [33], feedback vertex set [31, 30], and stable motifs [34], where each tactic provides a complimentary approach depending on the information available. We then sought to understand the impact of PC’s four most common gene mutations: KRAS (gain-of-function), TP53 (loss-of-function), SMAD4 (loss-of-function), and CDKN2A (loss-of-function) [14, 27, 28, 23]. For ease of notation, we elected to use the abbreviations KRAS (K), TP53 (T), CycD (C), and SMAD (S). Each of these mutations can be computationally achieved by a functional knock-in or knock-out command, that permanently turns the given Boolean function ON/OFF. Detailed tutorials are included in our data repository listed under Section 7.

Within the various mutation combinations of the PC model, we used our stochastic simulator based on the SDDS framework (Section 2. 3) to derive aggressiveness scores from simulated long-term trajectory approximations (Section 10. 1) [23]. The estimates in Figure 3 were tracked using phenotype expressions from only the PCC because the PSC is not considered malignant. Results showed that certain mutation combinations may indeed be more aggressive than others. We then performed statistical analysis on clinical gene expression data and derived survival curves that corroborated estimated aggressiveness scores.

**Figure 3:**
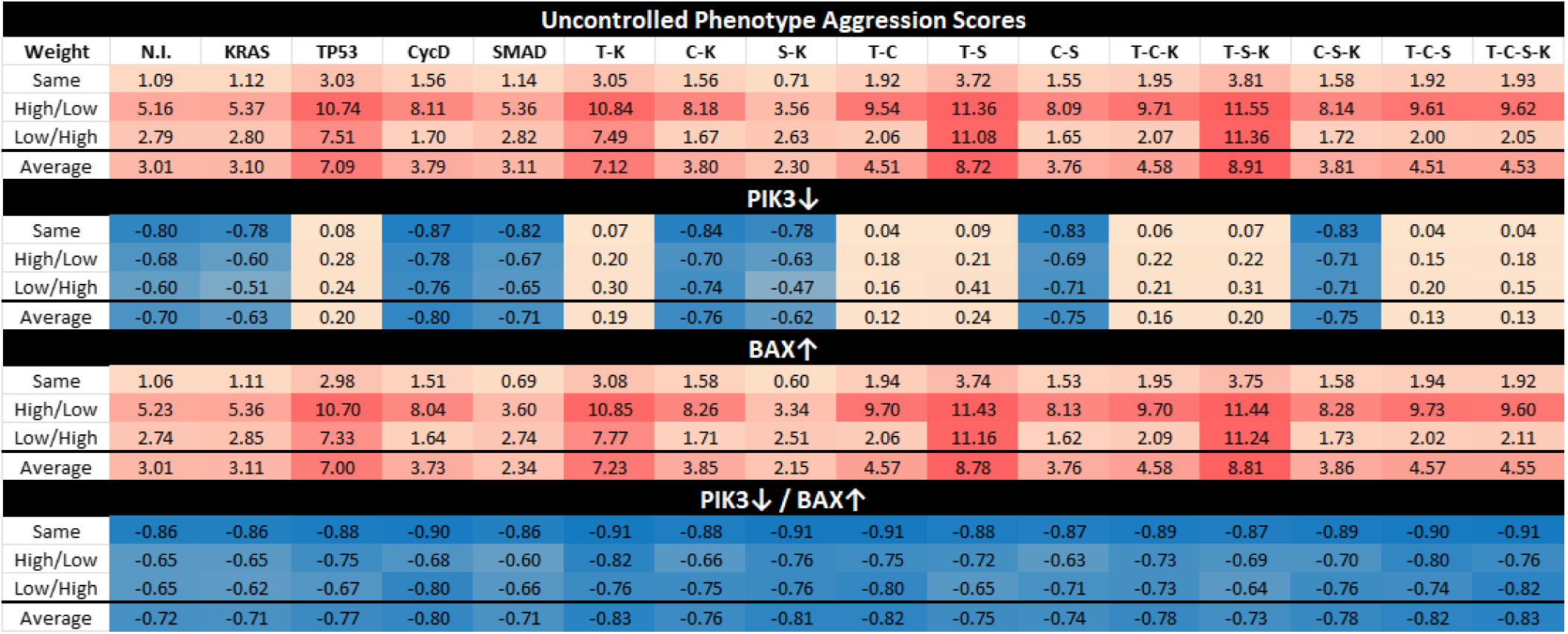
Aggression Scores and Target Efficacy. This figure shows aggression scores based on phenotype approximations by applying weights to trajectory probabilities. Here we include heat maps of aggression scores for each mutation combination (No mutation induced (N.I.), KRAS (K), TP53 (T), CycD (C), SMAD (S)), comparing cancer cell autophagy and proliferation while giving a negative weight to apoptosis. Row label “Same” indicates that the same weight was given to both autophagy and proliferation, “High/Low” indicates a high weight for autophagy but a low weight for proliferation, and “Low/High” indicates a low weight for autophagy but a high weight for proliferation. Scaling of the heat map ranges from orange (low score) to red (high score) based on the maximum and minimum values. Scaling shades of blue (cold) indicate non-aggressive or negative scores as a response to targets PIK3 knockdown (PIK3*↓*) and BAX agonist (BAX*↑*) [23]. See section 10. 1 for more details.

Phenotype control theory techniques revealed that sets of targets contained nodes within both the PCC and the PSC, highlighting PIK3 and BAX as a strong combination [23]. Notice that cells in Figure 10 contain duplicate pathways. While targets are found in both the PSC and the PCC, targets such as PIK3_c_ and PIK3_s_ can be considered as one target biologically. This is because a PIK3 inhibitor would act systemically rather than locally. Thus, when inducing controls in Figure 3, we assume systemic treatment. Here, we show a heatmap of projected mutation aggression compared to the application of target control. Note that as single agent targets, PIK3 knockdown (PIK3*↓*) and BAX agonist (BAX*↑*) are not universally effective. However, their combination is an effective control across all mutation combinations. We believe this can be explained by topological module rankings, detailed in Section 4. 1.

## 4 Results

The following key observations will be visited:

- Network “depth” of KRAS mutation can break the standard modular structure (4. 1)
- TP53 always directly impacts the module responsible for PCC apoptosis (4. 1)
- Certain modules may not determine phenotypic states (4. 1)
- Signaling from mutations to modules directly influences aggression (4. 2)
- Topological ranking gaps correspond to aggression projections (4. 2)
- Single agent targets are insufficient to out-compete downstream mutations (4. 3)

### 4. 1 Mutations perturb modular structure

In [1], the wild-type PC model from Figure 10 was first analyzed for modularity and revealed that the system contained three non-trivial modules. The module with top hierarchical rank in Figure 4, highlighted in yellow, is an autocrine loop of five nodes. Module 2, highlighted in green, contains thirty seven nodes spanning deep within both the PCC and PSC. The final module, highlighted in grey, is a negative feedback loop of only two nodes (referred throughout as the ‘duplex module’) [1].

**Figure 4:**
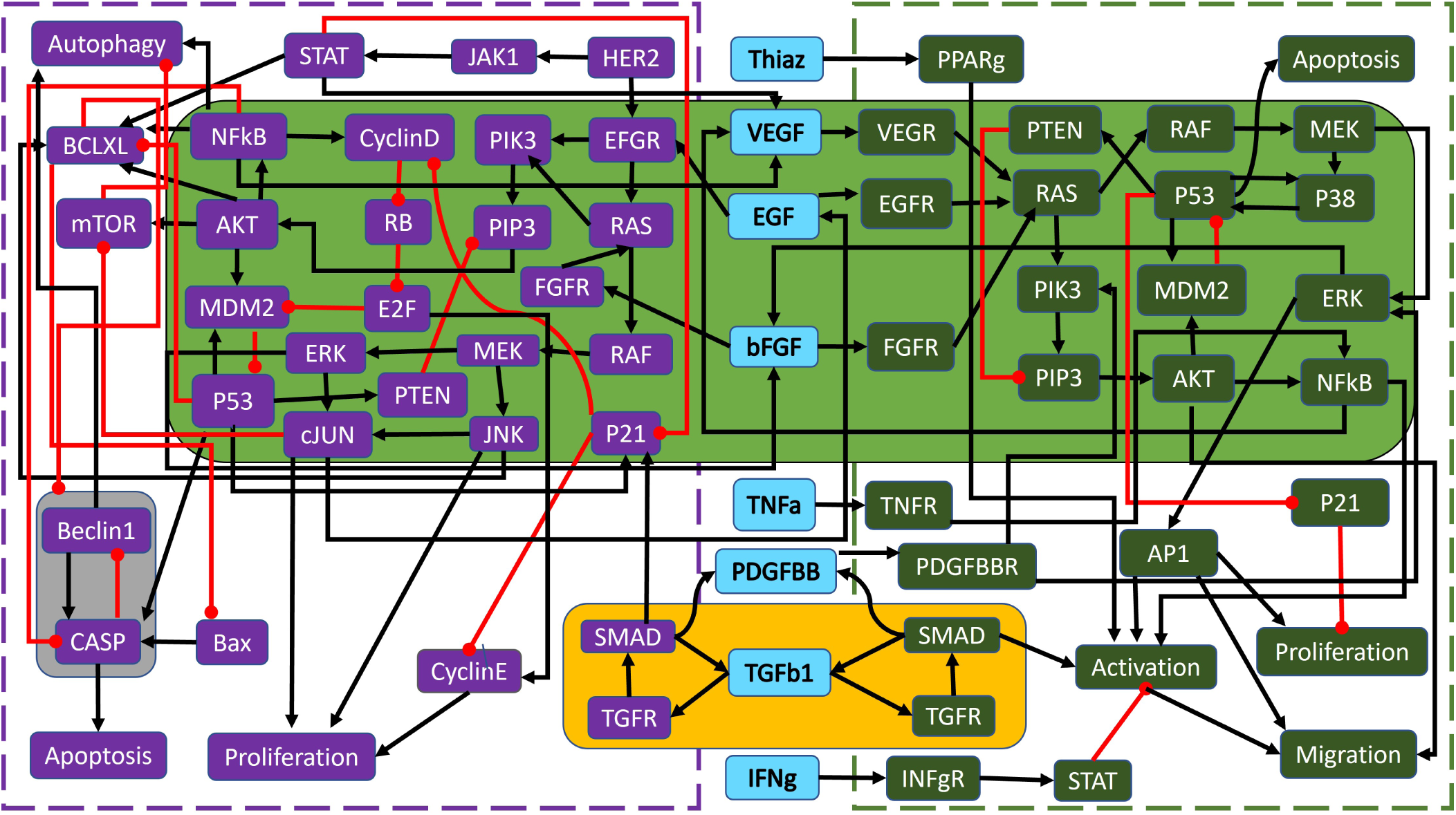
Wild-type PC wiring diagram with modules. Shown are the highlighted modules (yellow, green, and grey) for the wild-type PC model in Figure 10, adapted from [1]. Black barbed arrows indicate signal expression, while red oval arrows indicate suppression. A simplified structure can be seen in Figure 5b.

The overall wild-type modular structure from Figure 4 was achieved using the graphs in Figure 5. Notice that Figure 5a contains *all* modules, both trivial and non-trivial. However, Figure 5b is a reduced modular structure with only non-trivial nodes, including the cardinality of the modules as well as its original numbering from the condensation graph in curly brackets for easy reference. Figure 5a shows how phenotypes typically lie at the end (i.e. the bottom of a basin) because they are the dynamical endpoint: Apoptosis_s_ (24), Proliferation_s_ (26), Migration_s_ (28), Activation_s_ (27), Autophagy_c_ (21), Apoptosis_c_ (20), and Proliferation_c_ (23). Thus, the condensation ordering begins with source (or input) nodes and expands distally to phenotypes because the ultimate flow of the multicellular network goes from the middle outward. We anticipate that, due to this structure, we must use targets sufficiently upstream of phenotypes to subvert the impact of mutations.

**Figure 5:**
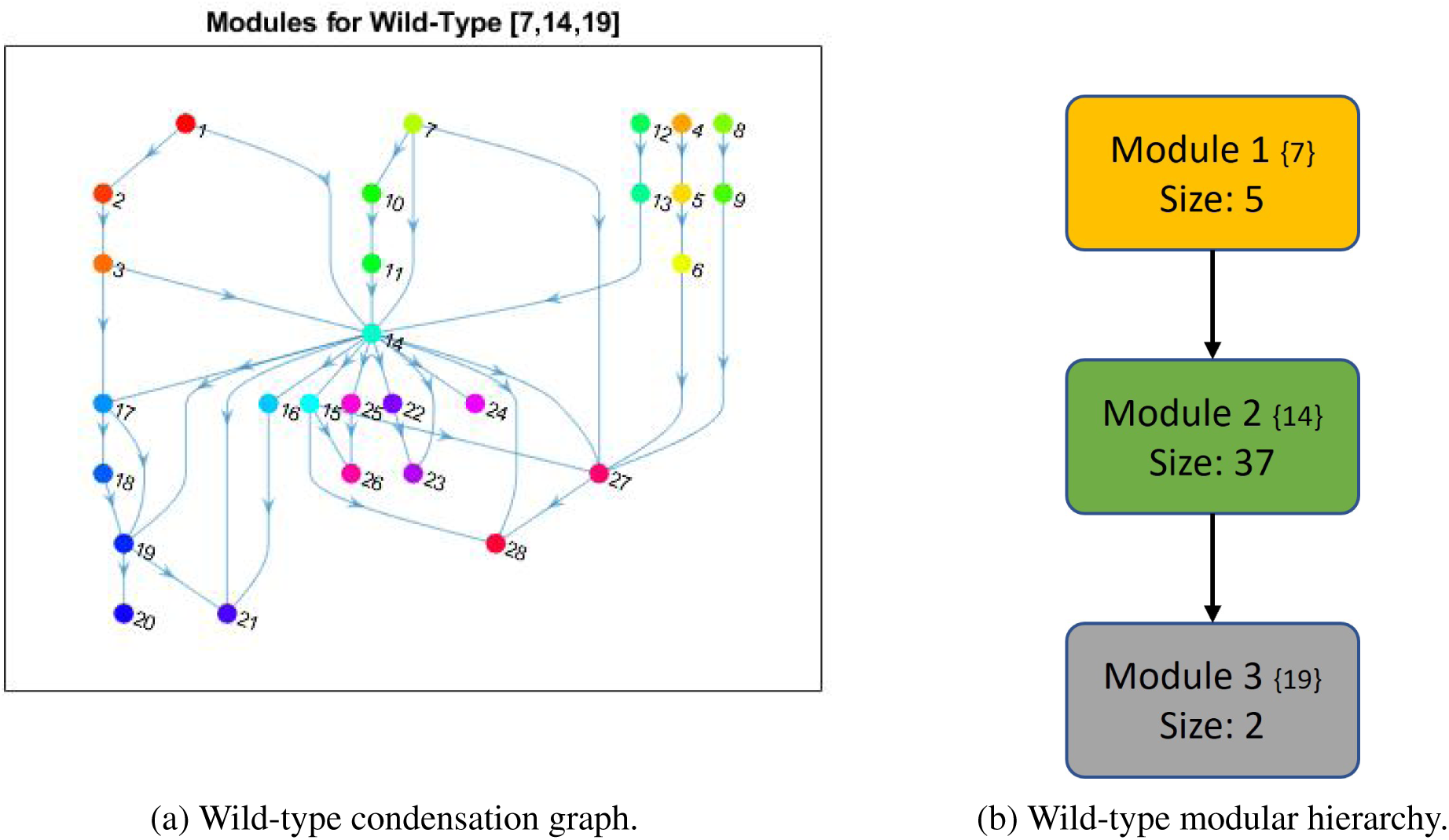
Wild-type PC modular structure. (a) Shown are the cumulative modules (trivial and nontrivial) ranked according to condensation. Node numbers are assigned by the MATLAB [32] “conncomp” function that bins connected graph components. (b) The reduced modular structure is shown with only nontrivial nodes, including the cardinality of the modules (Size) and its original condensation numbering in brackets (*{}*).

To better understand the impact of mutations on the modular structure, we constructed modules in same manner as the wild-type after mutations were induced. We give a comprehensive breakdown of module structures for each mutation combination in Figure 6. For example, to create Figure 6b, we first produced the corresponding condensation graph (Figure 12b) as previously described. We then identified the bins for each SCC, and noted direct or indirect communications between each module. Lastly, we identified location and influence of mutations that are indicated in red with respect to each module. We also included the PIK3 (P*↓*) / BAX (B*↑*) targets, indicated in blue with their respective locations and influences, to show the explicit topological impact of the intervention targets. Notably, mutations break SCCs and change the modularity structure both in magnitude of the modules (total nodes within a module) and amount of modules. More details with tables, graphs, files, and tutorials are available in the Supplementary materials and data repository (Sections 7 and 10).

**Figure 6:**
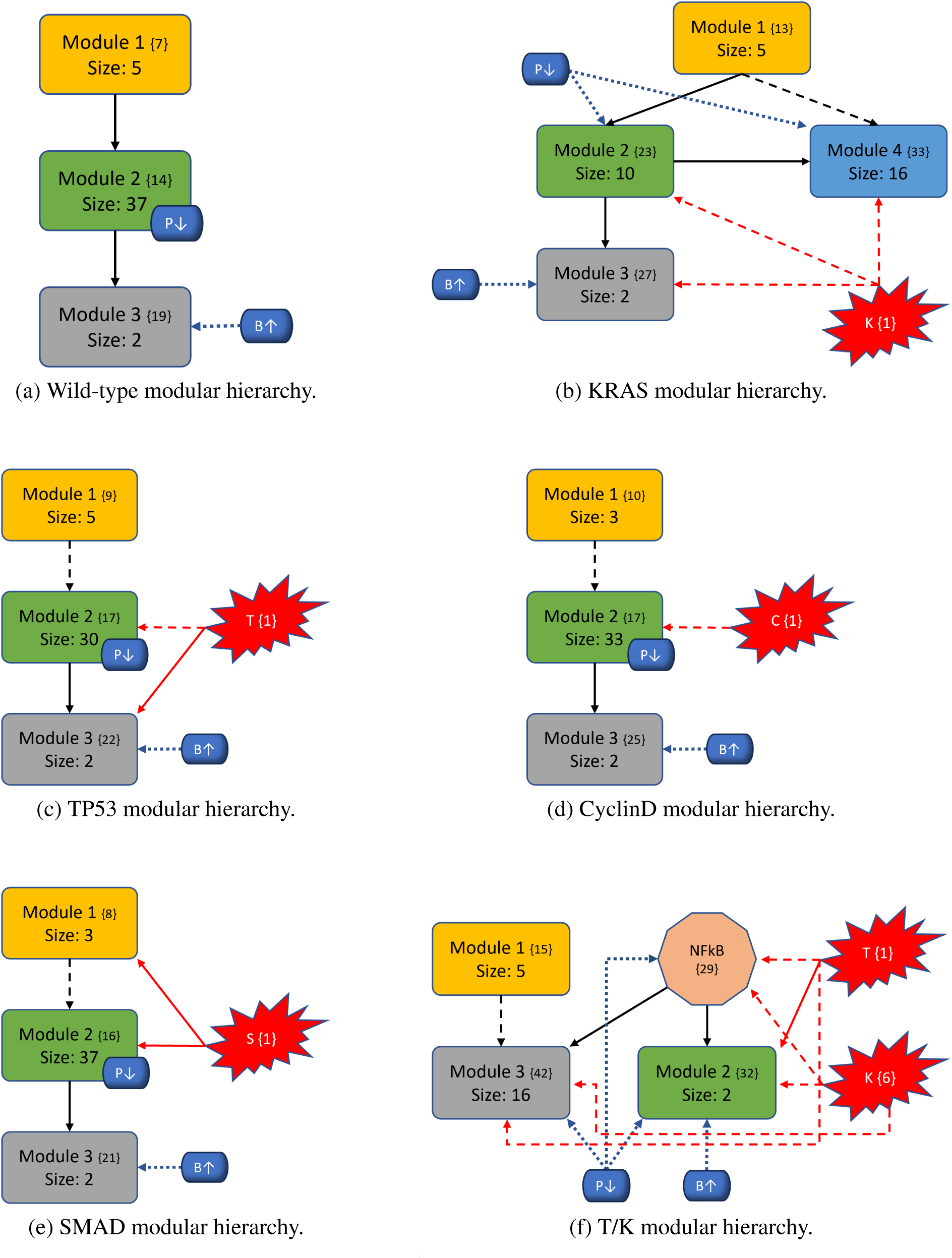
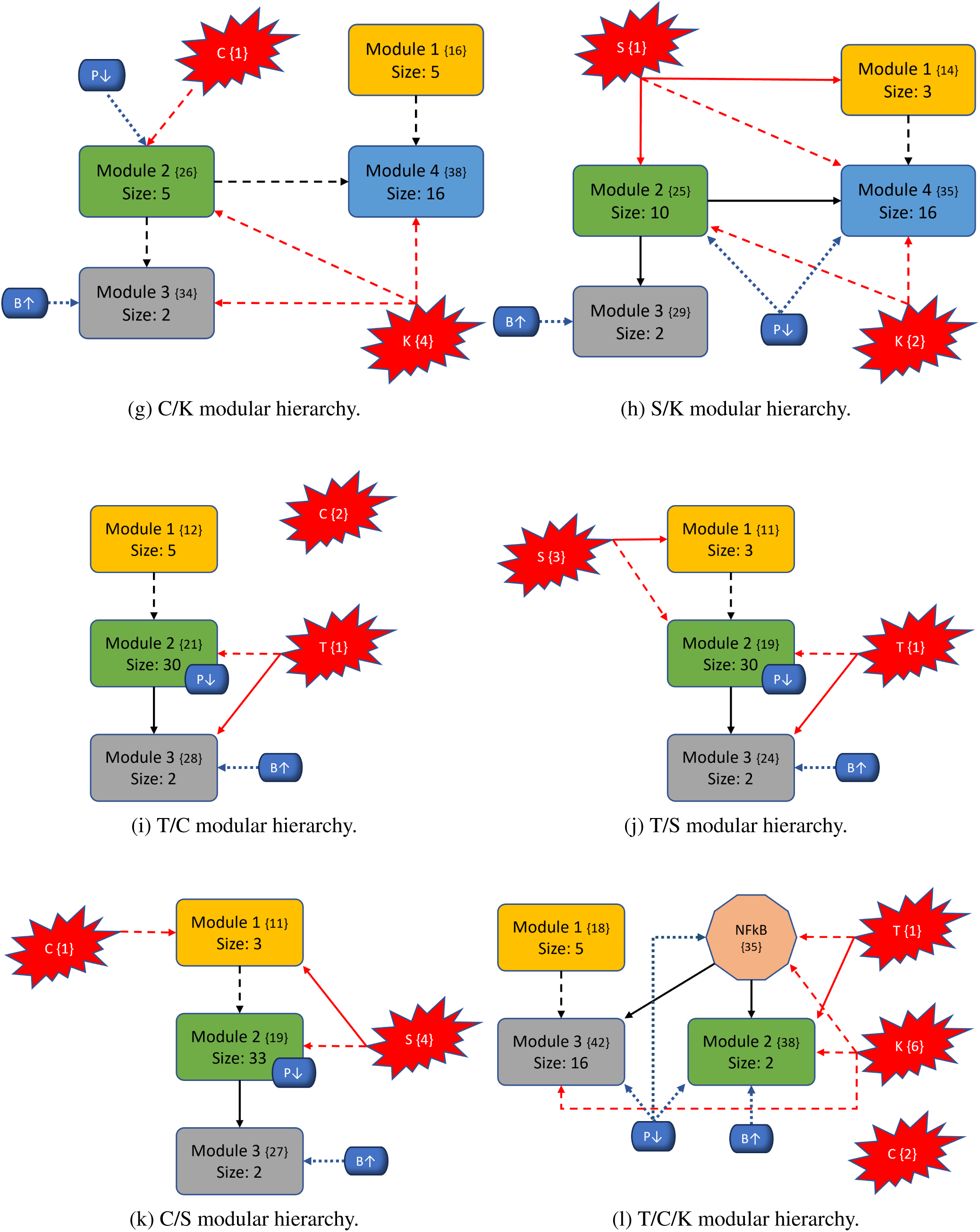
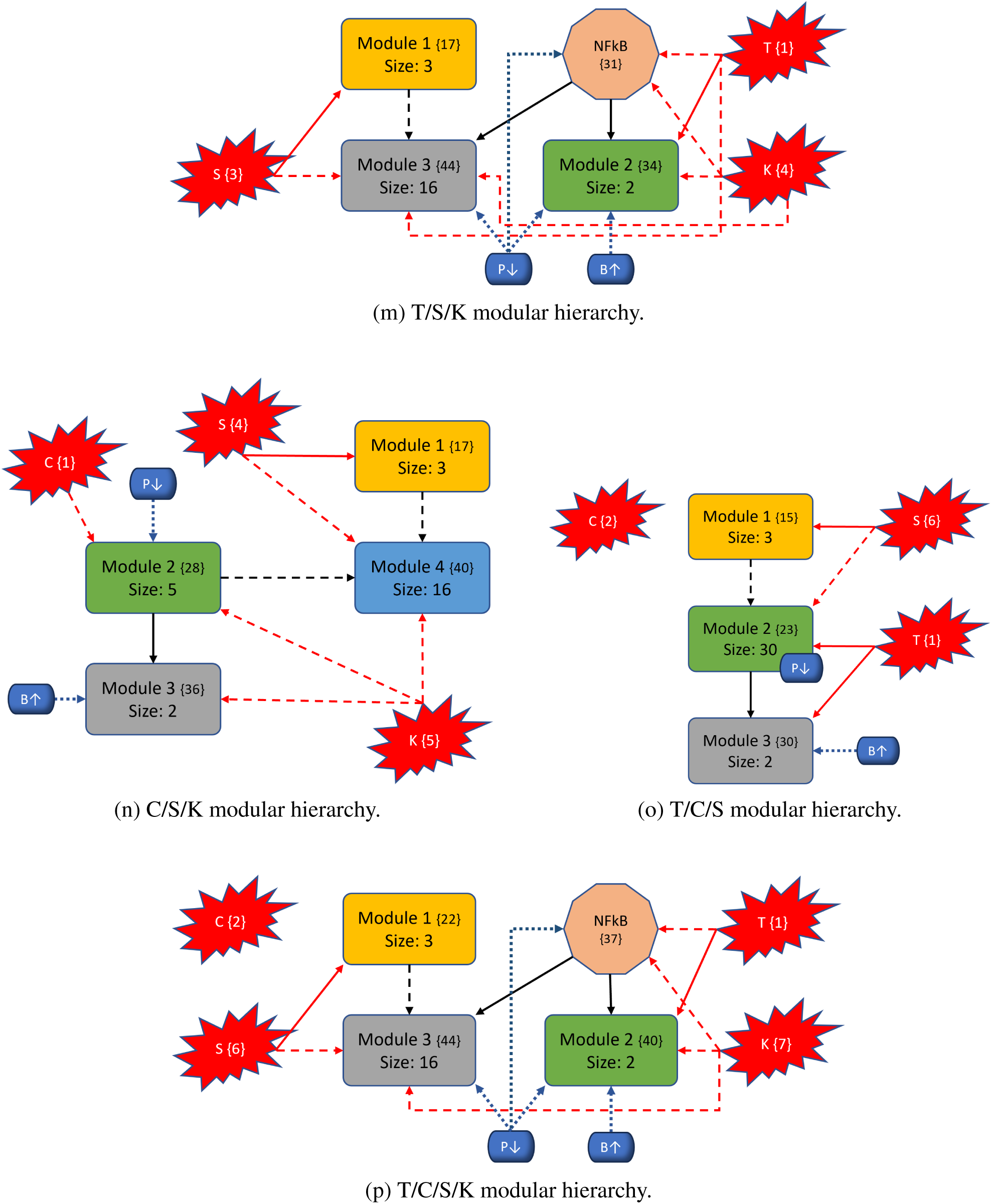
Variants of PC modular structure. Mutations are depicted with red “explosions” with their original module (condensation) numbers in brackets. Black arrows indicate communication between modules, red arrows indicate direct influence from mutations, and dashed arrows indicate indirect communication. Modules are ranked in order, and curly brackets contain the original numeration from the condensation graph. Lastly, the PIK3 (P*↓*) / BAX (B*↑*) targets are shown in blue with respect to their location inside or outside modules to show their downstream modular effects.

In general, we maintain the three basic structures shown in Figures 6a, 6b, and 6f. Some cases hold to the same structure as the wild-type, but notably, *every* instance of KRAS induction yields split modules - sometimes trivial (octagonal shape in Figure 6f) and other times non-trivial (separate blue node). The former observation of a nontrivial node split arises when KRAS is combined with TP53. We posit that these phenomena are due to the location of KRAS in regard to its “depth” within the largest central module. Notice that Figure 4 shows SMAD, TP53, and CyclinD are all located distally within their associated modules (Module 1 and all others in Module 2, respectively). Therefore, inducing their mutation merely shrinks the module size. When KRAS is induced, it is located more centrally within its original module and results in a broken component. For example, Figure 6b shows a gained module that only contains elements from the PSC that have split from their strongly connected PCC counterparts. Additionally, mutations do not always influence modules directly (e.g. Figure 6c), and there are scenarios where the mutation does not impact any modules at all (e.g. Figure 6o). Mutations yield anywhere from 28 to 51 modules in total (including trivial nodes), but non-trivial modules remain at either three or four in total (Figures 11a and 11b in Section 10. 3).

### 4. 2 Modular structure correlates with aggression

The most aggressive scores from Figure 3 are TP53, T/K, T/S, and T/S/K. The modularity structures for TP53 (Figure 6c) and T/S (Figure 6j) are unique, in that, the TP53 component always directly interacts with the duplex module that is responsible for apoptosis. Further, the structures for T/K (Figure 6f) and T/S/K (Figure 6m) are unique because (1) the duplex module is isolated from the standard downstream flow, and (2) the TP53 component influences all three downstream modules (versus only the trivial and duplex modules in T/C/K and T/C/S/K). This added layer of influence is what we believe drives a more aggressive prediction.

Figure 7 indicates the topological ranking for each module across every mutation combination using ‘toposort’. Rankings are calculated according to the modules overall percentile score (e.g. a rank of 5 among 28 total modules yields 18%). Thus, rankings of a higher percentile would indicate a greater impact (or stronger influence) on final phenotypic state. Figures 8 and 9 show the importance of module rankings and the gaps (distance) between them.

**Figure 7:**
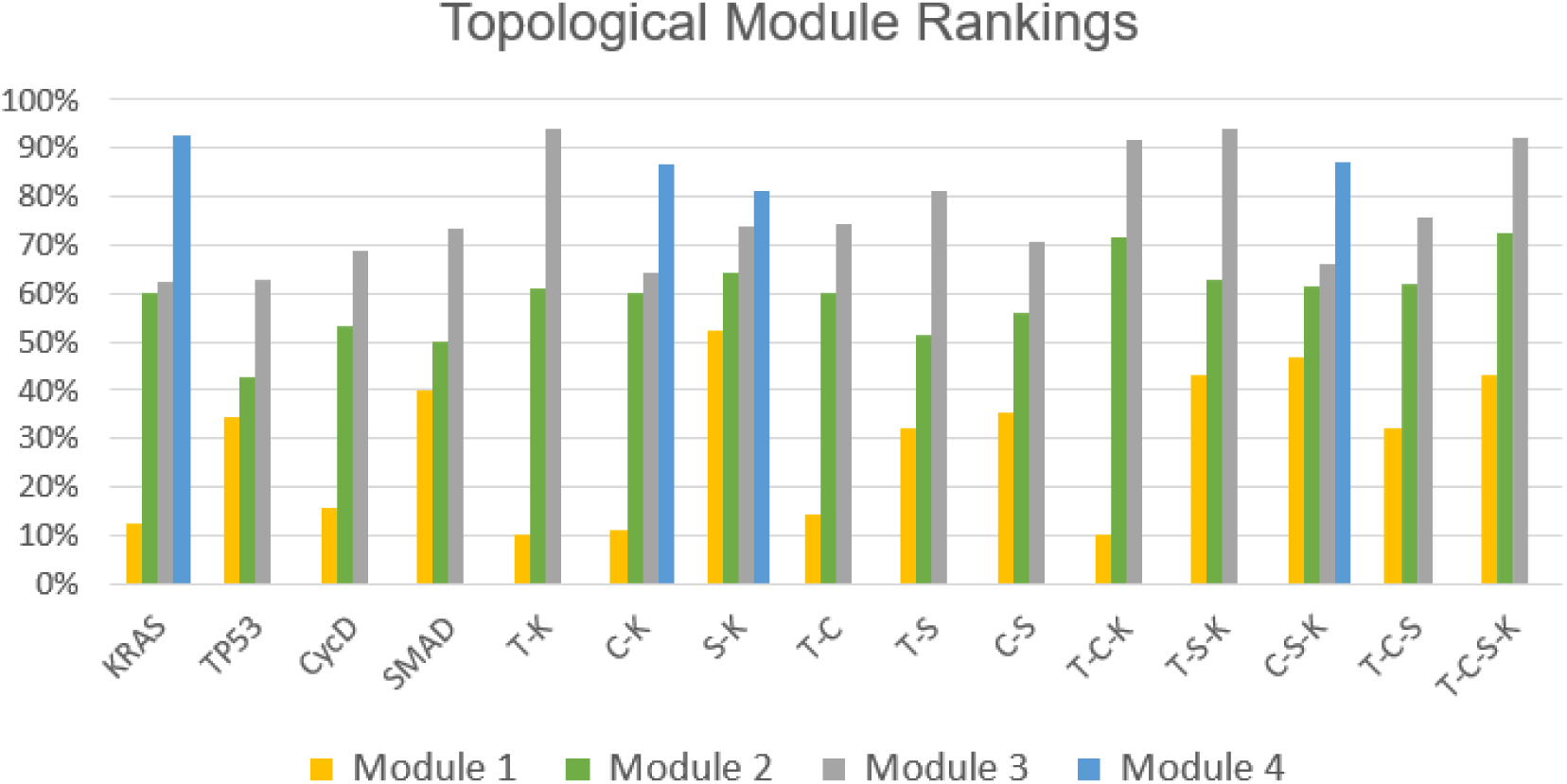
Toposort analysis. Shown are the overall topological rankings for each module among all mutation combinations. A table version is included on the data repository.

**Figure 8:**
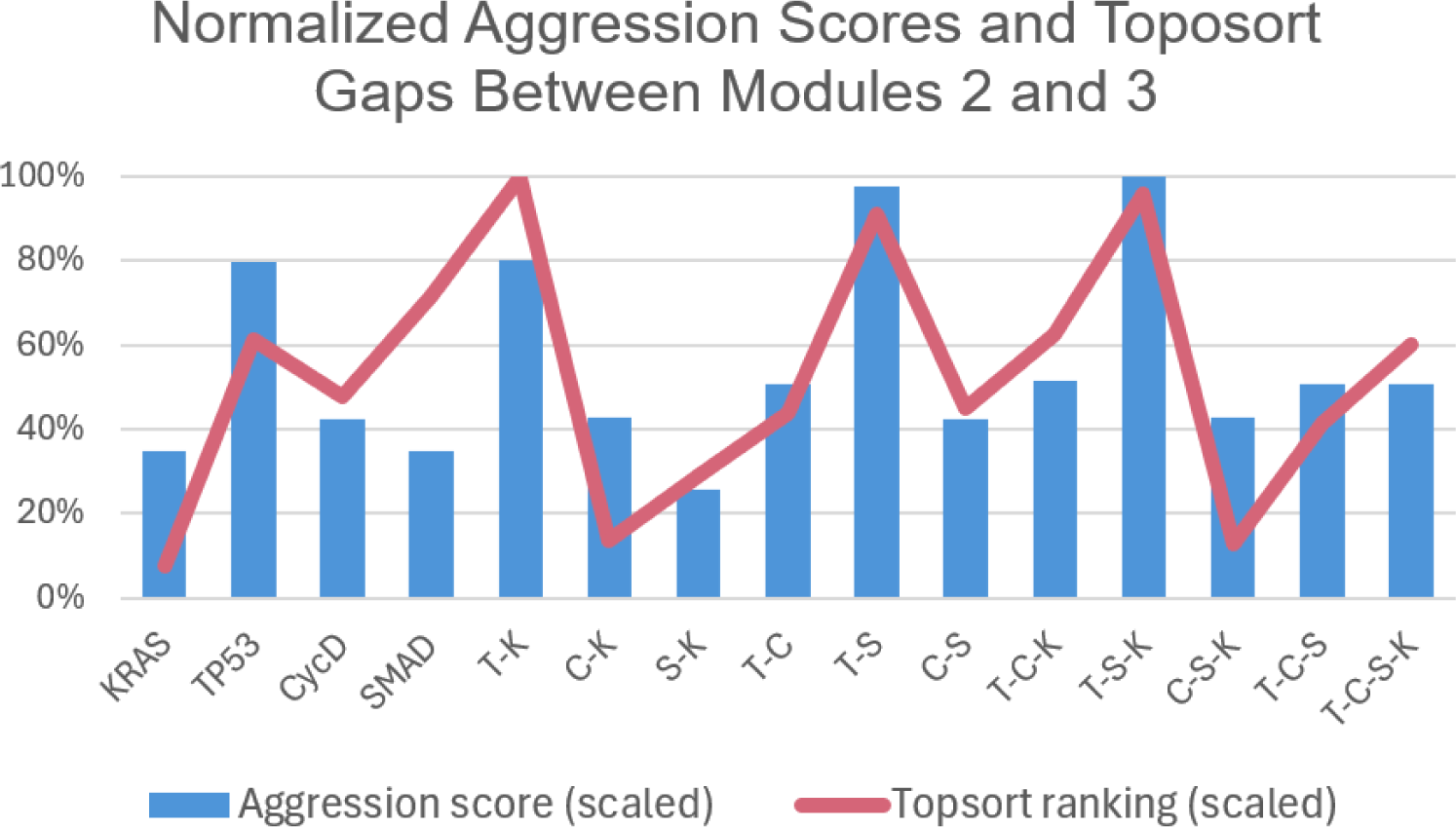
Normalized toposort gaps between Modules 2 and 3. After comparing the toposort rankings and gaps across all modules (Figure 7), the gaps between Modules 2 and 3 (i.e. the modules communicating to the PCC phenotypes) were identified as the most critical. Here we show a normalized scoring that indicates the largest variance (or distance) between Modules 2 and 3 best aligns with the predicted aggression scores from Figure 3.

**Figure 9:**
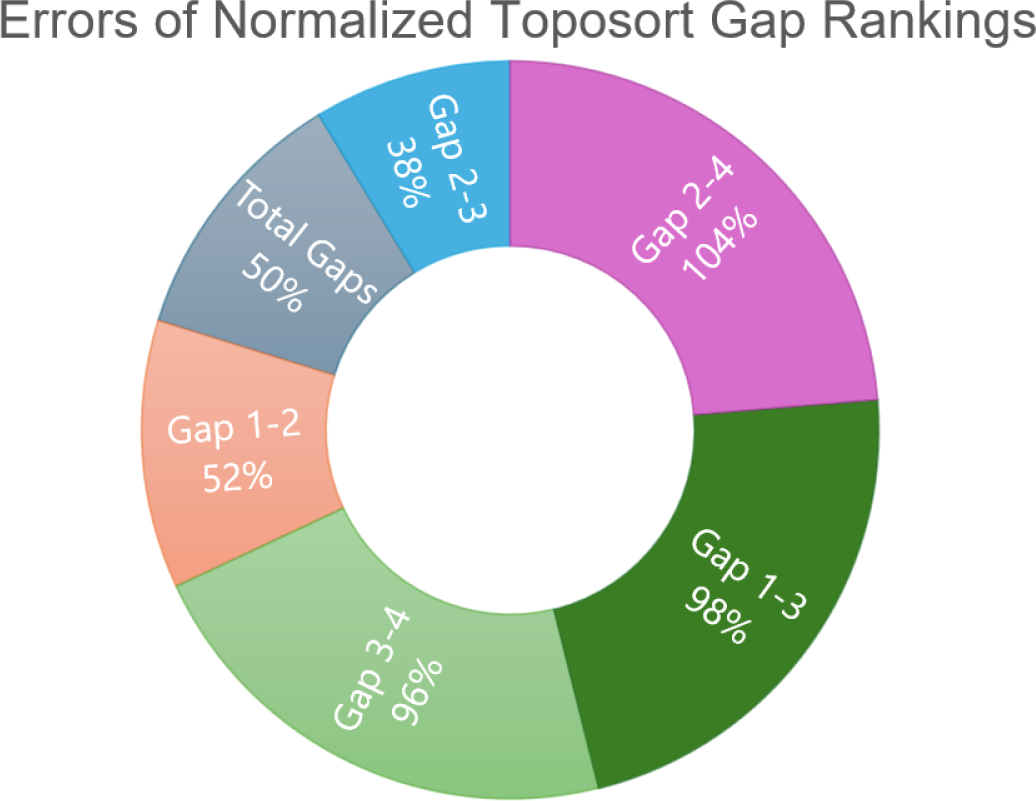
Errors of normalized toposort gap rankings. This chart is a comparison of average errors after ranking and normalizing toposort gaps, compared to the normalized aggression scores from Figure 3. Gaps between Modules 2 and 3 best explain the predicted aggression scores, further shown in Figure 8.

A compelling correlation is displayed in Figure 8, which shows the normalized percentage difference between the rankings of Modules 2 and 3, compared to previously predicted aggression scores. We posit that a greater distance between the most influential modules indicates a more difficult system dynamic to overcome when attempting to apply phenotype control theory. This may be attributed to more competing signals to overcome when compared to a small gap between modules. That is, gaps leave more room for noise, and therefore, link the topological structure to dynamical outcomes. This is further evidenced by comparing the average errors of all normalized module gaps, shown in Figure 9, where Gap 2-3 has the best fit to the aggression data. Indeed, greater gaps between these modules align with more aggressive scores.

### 4. 3 Mutations impact target efficacy

In [1], target analysis was performed on the wild-type PC model in Figure 4. For an overview of some of the most prominent control techniques, see [21]. Using apoptosis as the desired attractor, one could control the entire system by regulating a total of three nodes within the top two modules. The module with top hierarchical rank (yellow) is an autocrine loop with two fixed points and two 3-cycles. Therefore, the Feedback Vertex Set [30] strategy yielded control of Module 1 by pinning TGF*β*1. Next, Module 2 (green) was searched for targets using algebraic methods [33] and revealed that pinning KRAS in both the PCC and PSC stabilized the module. The final module (grey) becomes constant after the initial three pinnings. That is, the entire wild-type system was found to be controllable by a single growth factor and systemic KRAS inhibition, both within the first two modules [1].

However, KRAS is well-known for being unmanageable even though progress is being made towards its targetability [35]. Therefore, targeting PIK3 and BAX may be more achievable [36, 37, 23]. We showed in Figure 3 that the single agent controls of PIK3 and BAX are insufficient to achieve PCC apoptosis across all mutation combinations. This is because TP53 directly influences the duplex module responsible for apoptosis signaling (Figure 6) Since PIK3 is upstream of TP53, it must have help to circumvent or out-compete the mutation (through help from BAX). Such a combination is effective topologically because PIK3 is heavily involved in communications between multiple modules (either directly inside influential modules or indirect communication to many modules), and BAX can override the deepest mutation signals.

## 5 Conclusion

Systems biology is continually searching for general principles and tools for their identification. The approach of network modularity adds to the repertoire of techniques that provide structural and dynamical analysis across complex biological systems. We have shown here that the given definition of modules captures the drastic perturbing effect that mutational occurrence can have on normal biological mechanisms. A major limitation of most other approaches to modularity is their focus on a static representation of GRNs. Clearly, living organisms are dynamic and need to be modeled as such [1, 38]. That is precisely what we advocate through the framework established in Section 2. 1.

However, it is important to note that the decomposition presented in [1] does not preclude the existence of emergent properties. Each module is a complex dynamical system in itself. As we have shown, modules perturb other downstream modules, and their emergent properties propagate to other modules. That is, within each dynamical sub-system (module), the occurrence of mutations will have a direct and down-stream effect. Likewise, as the dynamics of upstream modules vary through time, the downstream modules may see an emergence of altered dynamical properties.

What we have provided herein uses a novel strategy of network modularity [1] to investigate the link between structure and function, using a previously published pancreatic cancer network as a case study. Namely, this link was shown to be strongly related to the variance in topological rankings of the most phenotypically influential modules. We have identified that the location of mutations, with respect to network depth and module positioning, expressly influence aggression and controllability. Thereby, presenting evidence that the impact and location of mutations with respect to modular structure directly corresponds to the efficacy of single agent treatments *in silico*. These cumulative results help provide more clarity on the impact of mutations in PC and posit the viability of using network modularity to study dynamical systems.

## 6 Acknowledgments

DP was supported by the NIH Training Grant T32CA165990. DM was partially supported by a Collaboration grant (850896) from the Simons Foundation.

## 7 Data Availability

All data and code used for running simulations, statistical analysis, and plotting is available on a GitHub repository at https://github.com/drplaugher/PCC_Mutations. The repository also includes “How-To” documentations for reproducibility.

## 8 Author Contributions

Conceptualisation: DP, Methodology: DP and DM, Software: DP, Formal analysis: DP, Supervision: DM, Writing—original draft: DP, Writing—review and editing: DP and DM

## 10 Supplementary material

### 10. 1 Aggression scores

We derived aggressiveness scores for each mutation combination using long-term trajectory approximations. Simulations were run using 1000 random initializations, 300 time steps, and 1% noise to achieve an approximate probability of phenotype expression. In Table, 1, we see that the non-induced (N.I.) system showed levels of 51% autophagy, 36% apoptosis, and 21% proliferation. The heat maps in Figure 3 are sorted with column-wise mutation groups and used to compare cancer cell autophagy and proliferation while giving a negative weight (*ω* = *−*1) to apoptosis. The row label “Same” indicates that the same weight was given to both autophagy and proliferation (used value *ω* = 2 for both), “High/Low” indicates a high weight for autophagy (*ω* = 10) but a low weight for proliferation (*ω* = 2), and “Low/High” indicates a low weight for autophagy (*ω* = 2) but a high weight for proliferation (*ω* = 10). Thus, scores were calculated using:

**Table 1:** Expression Approximations. This table records the approximate phenotype expressions for the PCC in Figure 10. Given 1,000 random initial states, these results show trajectory approximations after 300 time steps (i.e. function updates) with 1% noise.

|  | Uncontrolled Trajectory Approximations |  |  |  |  |  |  |  |  |  |  |  |  |  |  |  |
| --- | --- | --- | --- | --- | --- | --- | --- | --- | --- | --- | --- | --- | --- | --- | --- | --- |
|  | N.I. | KRAS | TP53 | CycD | SMAD | T-K | C-K | S-K | T-C | T-S | C-S | T-C-K | T-S-K | C-S-K | T-C-S | T-C-S-K |
| Aut<sub>c</sub> | 0.509 | 0.531 | 0.964 | 0.818 | 0.528 | 0.974 | 0.827 | 0.356 | 0.953 | 0.955 | 0.817 | 0.97 | 0.967 | 0.82 | 0.962 | 0.961 |
| Apo<sub>c</sub> | 0.359 | 0.356 | 0.021 | 0.109 | 0.343 | 0.011 | 0.118 | 0.479 | 0.025 | 0.035 | 0.105 | 0.018 | 0.013 | 0.1 | 0.028 | 0.017 |
| Pro<sub>c</sub> | 0.213 | 0.209 | 0.56 | 0.017 | 0.211 | 0.555 | 0.013 | 0.24 | 0.018 | 0.92 | 0.012 | 0.015 | 0.944 | 0.018 | 0.01 | 0.014 |

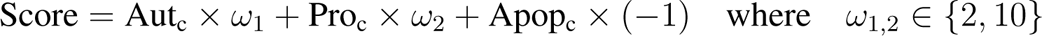

Scaling of the heat map ranges orange (low score) to red (high score) based on the maximum and minimum values in each table. However, blue shading (i.e. cold) indicates a negative score, which is interpreted as successful depletion of aggression. See [23] sections 2.3 and 4.4 for more details.

Lastly, we justify the positive weight given to autophagy, which is a natural process where cells heal themselves. The cell will break down any damaged or unnecessary components, and it will reallocate the nutrients from these processes to those that are essential. However, studies have shown that autophagy is required for pancreatic tumor growth [39]. Autophagy can help tumors overcome conditions such as hypoxia and nutrient deprivation. Within tumors, cells can exist under hypoxic conditions. If activated autophagy is then suppressed by deletion of Beclin 1, studies have shown increased cell death. It has also been observed that autophagy is increased in KRAS mutated cells, and aids in survival of the cancer cells while experiencing nutrient starvation. Further, animal studies have shown that autophagy contributes to tumor-cell survival by enhancing stress tolerance and supplying nutrients to meet the metabolic demands of tumors. Once suppression of autophagy occurred, there was an observance of tumor-cell death [40].

Note: our aggression scores are based on combinations autophagy, apoptosis, and proliferation, merely one method among many for estimating aggression. Moreover, the attractor analysis (see [11, 23] indicated that certain mutation combinations yield a large basin for attractors with both autophagy and proliferation expression. It is likely that modular structure alone is not enough to determine aggression and target cardinality. Rather, it should be used alongside other analyses.

### 10. 2 Boolean pancreatic cancer model and functions

**Table 2:**
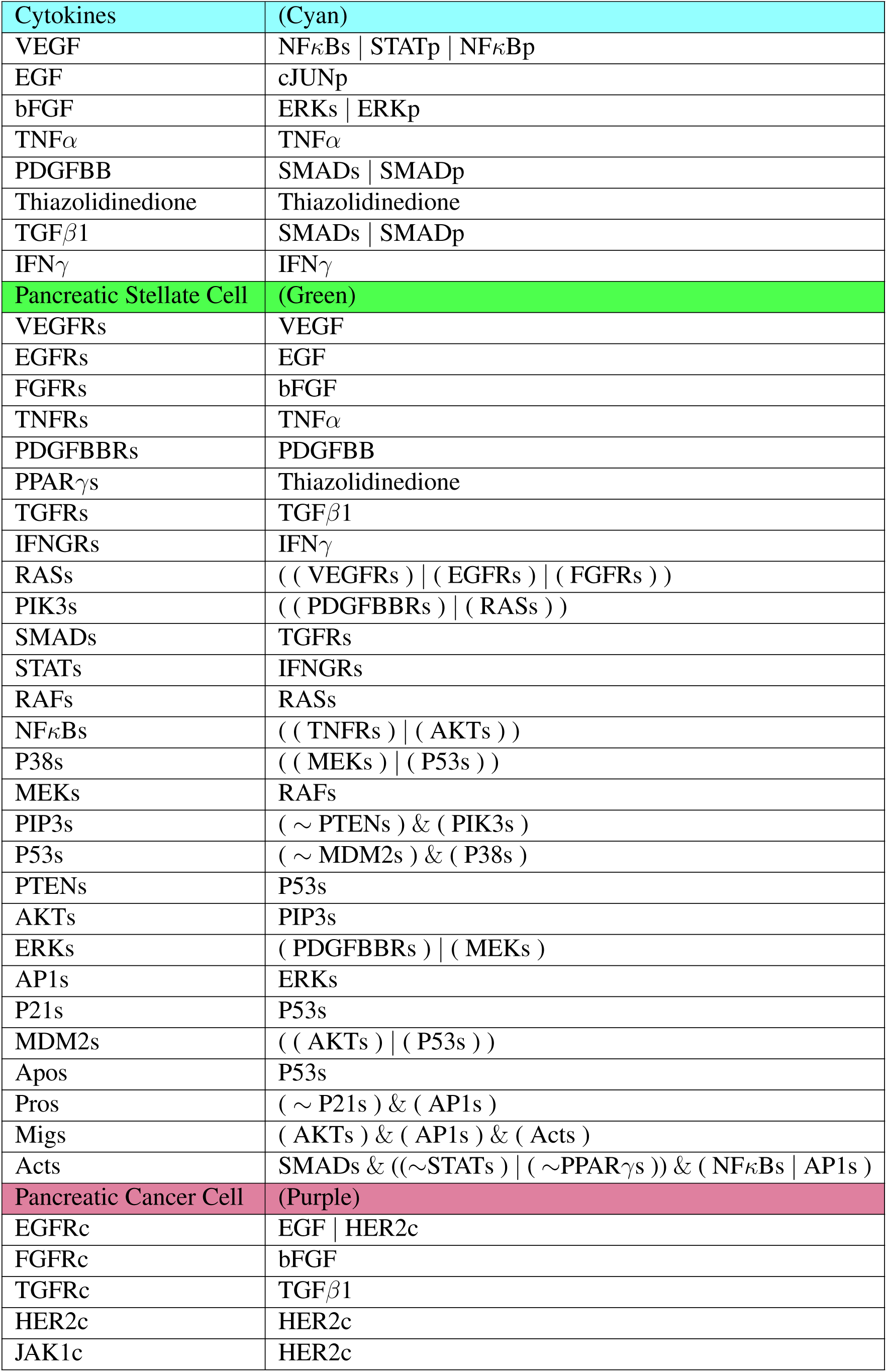
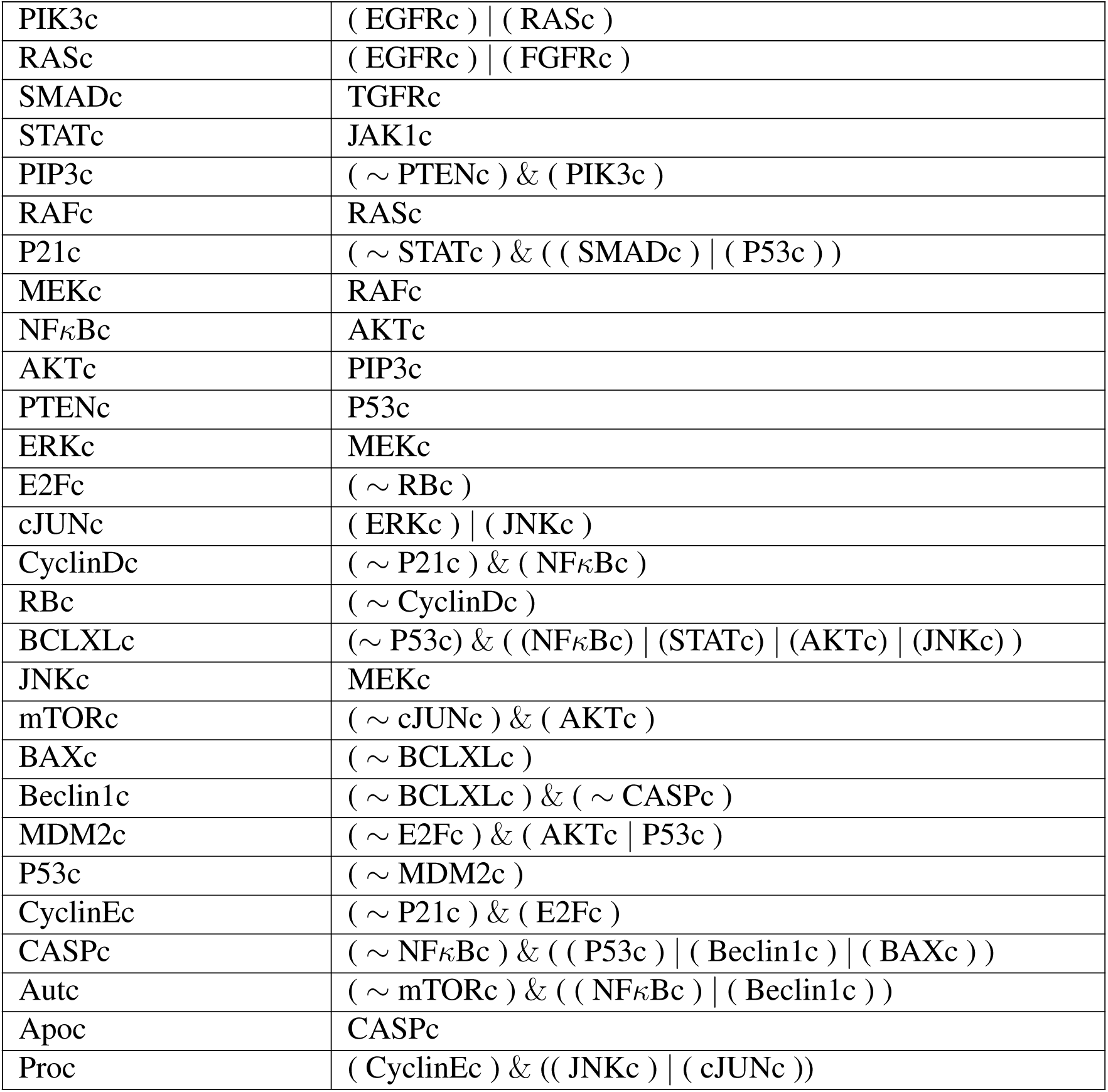
Boolean functions for the whole pancreatic cancer model. Each function indicates the next state of the node in terms of the current states of said nodes’ regulators. Activation is written as OR statements, while suppression is written as AND NOT. The exception to this rule is PCC proliferation, because of its upstream signaling.

**Table 3:**
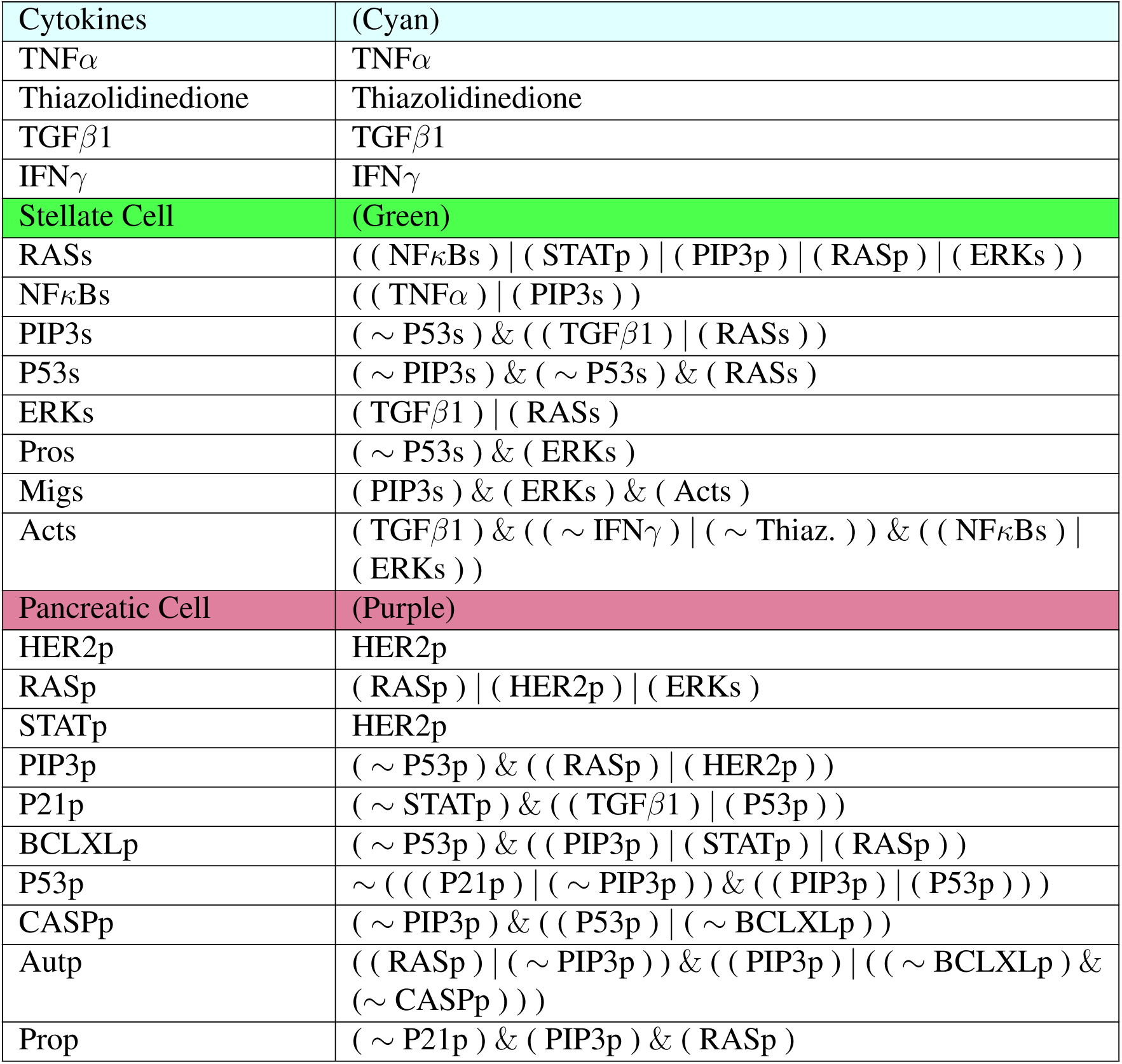
Boolean functions for the reduced pancreatic cancer model. Each function indicates the next state of the node in terms of the current states of said nodes’ regulators. Activation is written as OR statements, while suppression is written as AND NOT. Functions maintain the rules from the whole model by substituting values from the deleted nodes.

**Figure 10:**
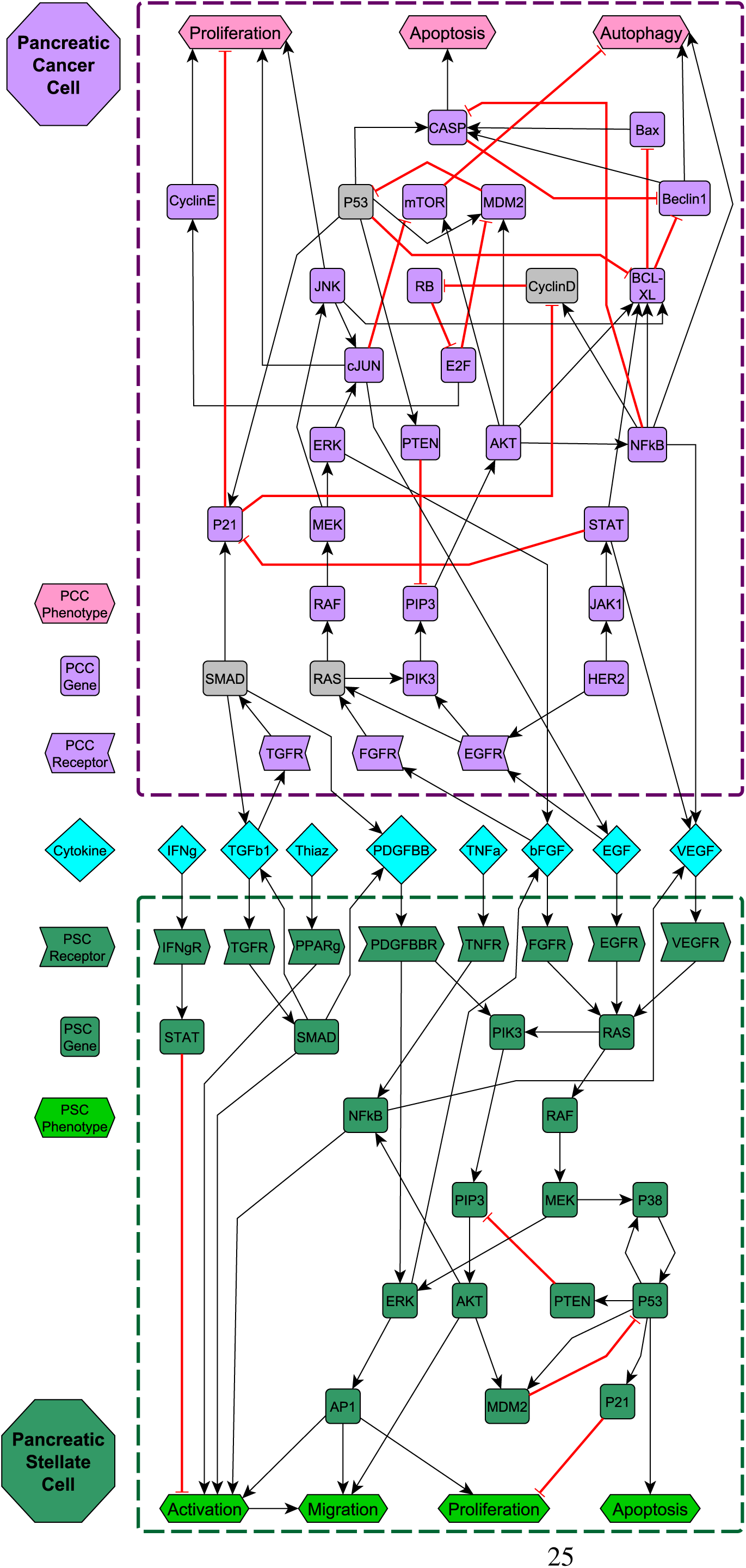
Gene regulatory network model of pancreatic cancer. Shapes and colors of nodes indicate their function and cell type (respectively), as shown in the legend. Black barbed arrows indicate signal expression, while red bar arrows indicate suppression. Grey nodes located in the PCC indicate prevalent mutant genes [11, 29].

### 10. 3 Tables and Graphs

**Figure 11:**
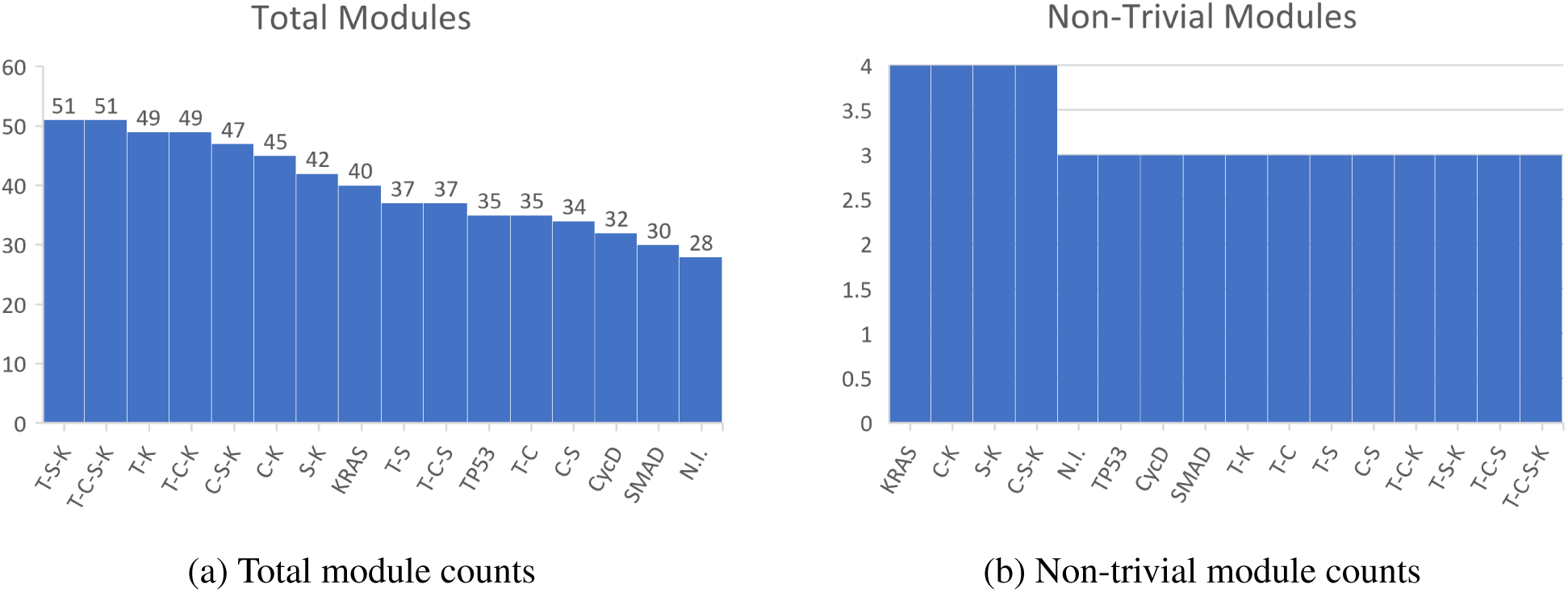
Module counts.

**Figure 12:**
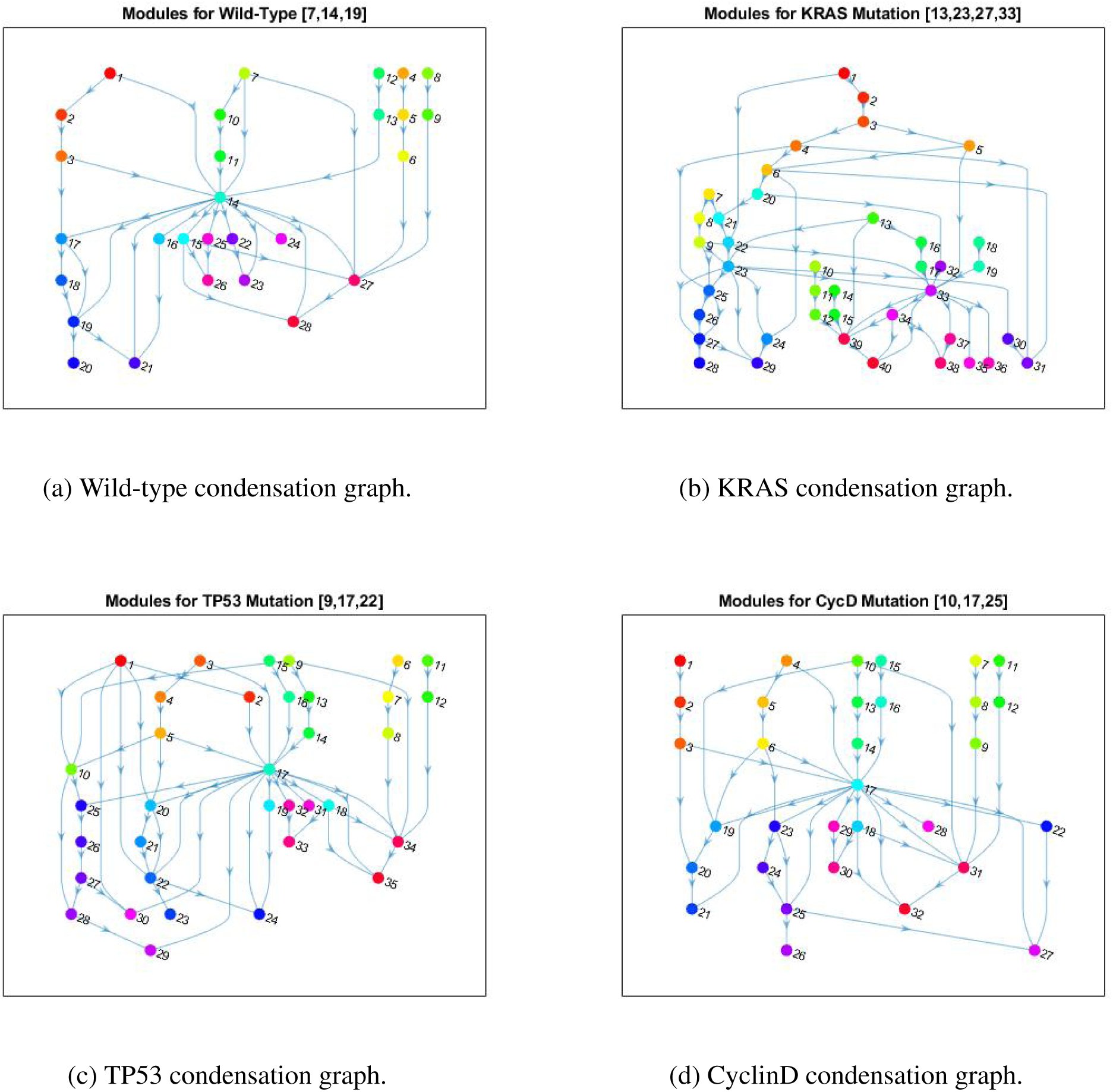
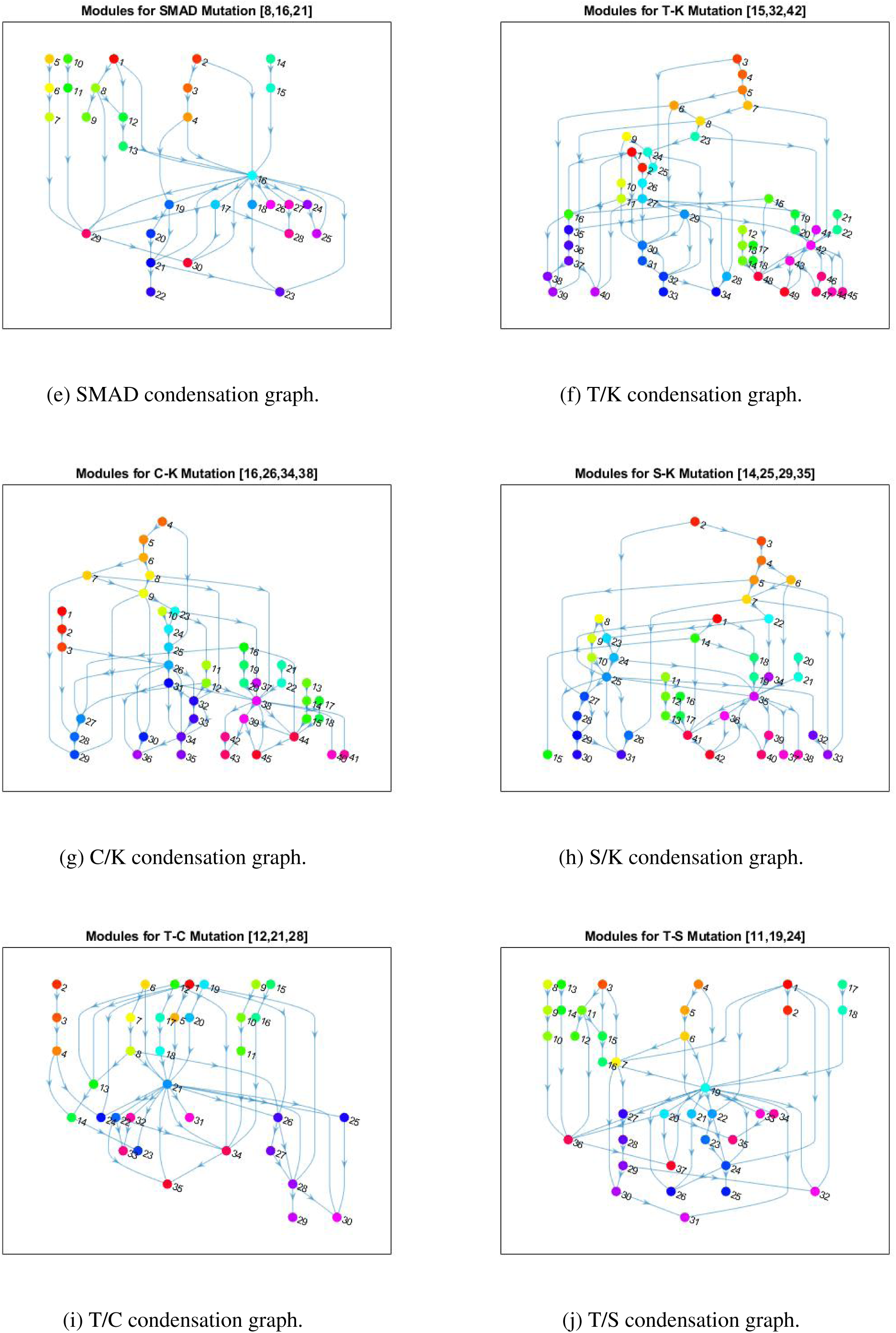
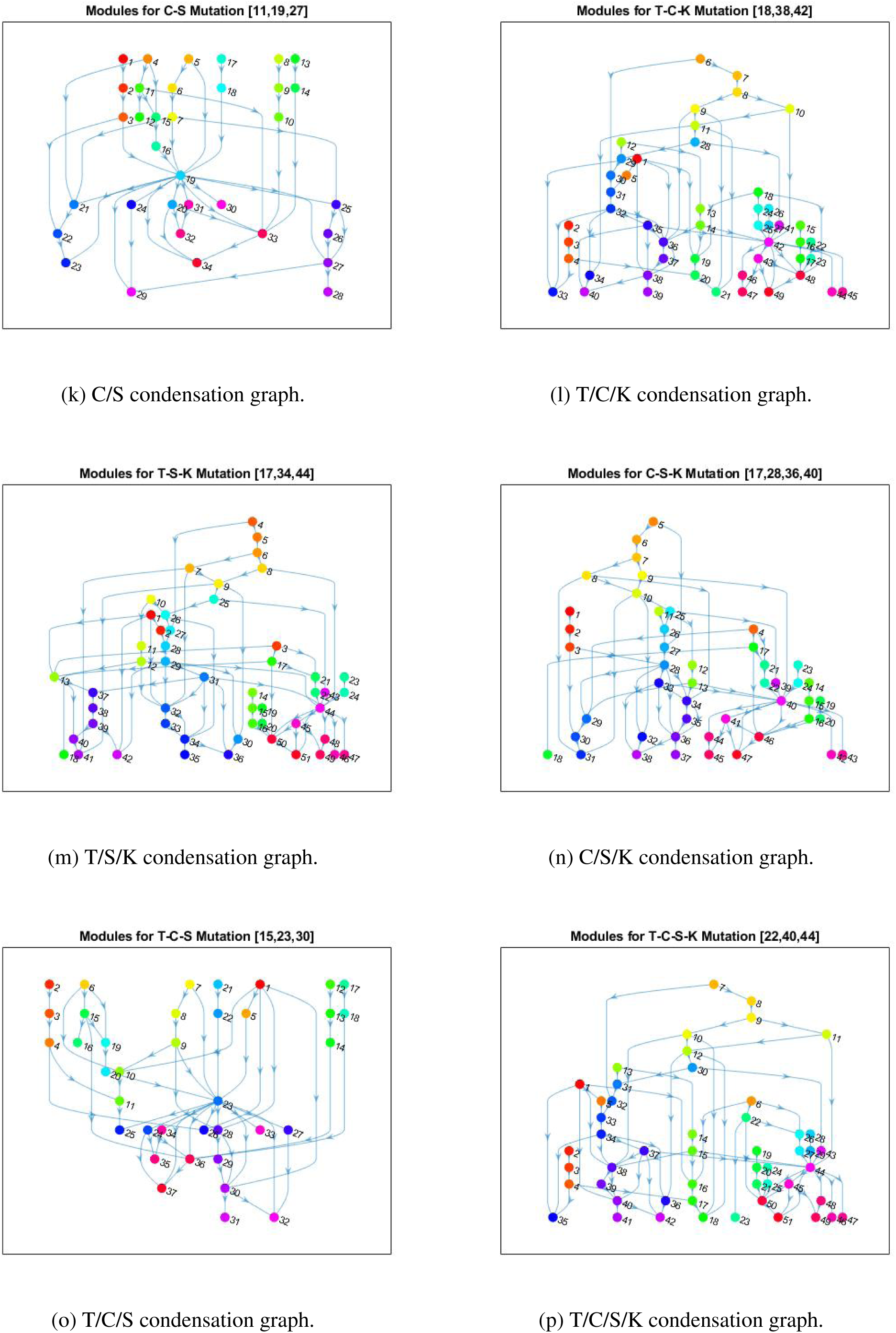
PC condensation graphs. Included are all condensation graphs for each mutation combination. These are directed, acyclic graphs that are topologically sorted, and whose nodes represent the strongly connected components of Figure 10. Colors of nodes are based on the components they represent, and node numbers correspond to bin numbers (see supplementary materials through Section 7).

